# Omura’s whale (*Balaenoptera omurai*) stranding on Qeshm Island, Iran: further evidence for a wide (sub)tropical distribution, including the Persian Gulf

**DOI:** 10.1101/042614

**Authors:** Sharif Ranjbar, Mohammad Sayed Dakhteh, Koen Van Waerebeek

## Abstract

A small, juvenile rorqual live-stranded on Qeshm Island, Iran, in the northern Strait of Hormuz (Persian Gulf) in September 2007. Cause of stranding remains unknown but the whale (QE22.09.2007) showed no severe traumatic injuries nor was emaciated. Based on at least seven morphological features, considered diagnostic in combination, allowed a positive identification as Omura’s whale *Balaenoptera omurai*. Features included diminutive body size (397 cm), a large number of ventral grooves (n=82) extending caudad of the umbilicus, a strongly falcate dorsal fin, asymmetric colouration of the head (especially lower jaws) reminiscent of fin whale, including three unilateral dark stripes, faint/incomplete lateral rostral ridges, record low number of short, broad baleen plates (204 in right jaw). The likelihood for the existence of a local *B. omurai* population in the eastern Persian Gulf or northern Arabian Sea seems higher than the wandering of a very young animal or mother/calf pair from any of the known distant distribution areas in the eastern Indian Ocean or SW Indian Ocean (Madagascar). This is the first record of *B. omurai* in the NW Indian Ocean.

## Introduction

A small rorqual (*Balaenoptera* sp.) with a body length of 397 cm stranded at N26°56’17.88”, E56°16’42.09” on Qeshm Island, Iran, in the northern Strait of Hormuz, Persian Gulf, on 22 September 2007. The diminutive rorqual, evidently a calf or juvenile, had stranded alive on 21 September 2007, at an indeterminate time. During the night a team of local volunteers attempted to rescue the animal by keeping it wet with the plan to refloat it with rising tide, actions which were filmed by Mr. Besharati Asghar. Predictably the rescue failed and the whale died *in situ*.

Based on external morphological characteristics we here identify the Qeshm rorqual as an Omura’s whale *Balaenoptera omurai* Wada, Oishi & Yamada 2003. This recently described, small rorqual species, with an adult size of barely 11.7m, was until 2015 thought to be distributed exclusively in tropical and subtropical waters of the southwestern Pacific and the eastern Indian Ocean (Wada et al. 2003; Jefferson et al. 2008; Reilly et al. 2008a). Cerchio et al. (2015) extended known range to the Southwestern Indian Ocean when reporting on a population resident in coastal waters of northern Madagascar. These authors documented external morphology in some detail from underwater photography, which resulted essential in the present analysis. Simultaneously, Jung et al. (2015) demonstrated at least occasional distribution of *B. omurai* into the Northeast Atlantic Ocean, following the genetic identification of a 398 cm juvenile stranded on an Mauritanian beach in 2013. Jung et al. (2015) advanced two hypotheses, either an unrecognised Atlantic population or, less likely, an inter-oceanic vagrant.

The main significance of the Qeshm specimen resides in the fact that *B. omurai* has not before been documented from the Persian Gulf, Iran and the Northwestern Indian Ocean. The possible occurrence off Iran was recognized by Braulik et al. (2010).

## Material and Methods

On 22 September, two of us (SR, MSD) examined, measured and photographed the fresh carcass, henceforth referred to as specimen QE22.09.2007. Multiple shallow cuts and abrasion injuries, apparently linked to the stranding event, were present but there was no indication of major trauma or emaciation. As the specimen was not necropsied and no tissue samples were collected, the cause of death will remain unknown. Considering that it was initially thought to be a Bryde’s whale, the carcass was buried on the beach for later retrieval but, washed out during a spring tide, it eventually was lost. Numerous photographs (n=124) and a video of both the live and freshly dead (<12 hours) whale were deposited at the Qeshm Environment Administration at Qeshm City, with back-up copies now archived at the Peruvian Centre for Cetacean Research, Lima.

As to positively identify QE22.09.2007 based on its distinctive morphology and pigmentation pattern, we followed an inferential process in which at least seven morphologic features described for *B. omurai* (Wada et al., 2003; Jefferson et al., 2008; Cerchio et al., 2015) were demonstrated to be present and, combined, deemed diagnostic for *B. omurai*. QE22.09.2007 was also distinguished from the other seven known rorqual species (genus *Balaenoptera* Lacépéde) in a differential diagnosis, discarding blue whale, *B. musculus* (Linnaeus, 1758); fin whale, *B. physalus* (Linnaeus, 1758); sei whale, *B. borealis* Lesson, 1828; Bryde’s whale, *B. brydei* Olsen, 1913; Eden’s whale, *B. edeni* Anderson, 1879; Antarctic minke whale, *B. bonaerensis* Burmeister, 1867; and common minke whale, *B. acutorostrata* Lacépéde, 1804.

Although increasing evidence (e.g. Wada et al., 2003; Sasaki et al., 2006) suggests that large-sized Bryde’s whales (*B. brydei* Olsen, 1913) form a clade distinct at specific level from the smaller-sized Eden’s whale (*B. edeni* Anderson, 1879), several authors continue to defend a conservative view of conspecificity where *B. brydei* is considered a junior synonym of *B. edeni* (e.g. Best, 2003; Reilly et al. 2008b). Although we tend to agree with dual species (Wada et al. 2003), in practical terms it remains difficult to assign individual specimens to *brydei* or *edeni* forms since most published variation (ranges) in external measurements, meristics and other phenotypic features span the pooled intraspecific variation of both large and small forms of Bryde’s whales *sensu lato*.

We counted the throat (ventral) grooves from high-resolution photographs at the cross-section of the pectoral fins over an ½ exposed throat (i.e. from the ventral midline up to one flipper). Next, we added the count of the few grooves lateral to one pectoral fin, visible from a side view photo. Throat grooves were counted several times independently, then averaged. Finally, we multiplied by 2 to provide the total number of throat grooves.

**Figure 1.**
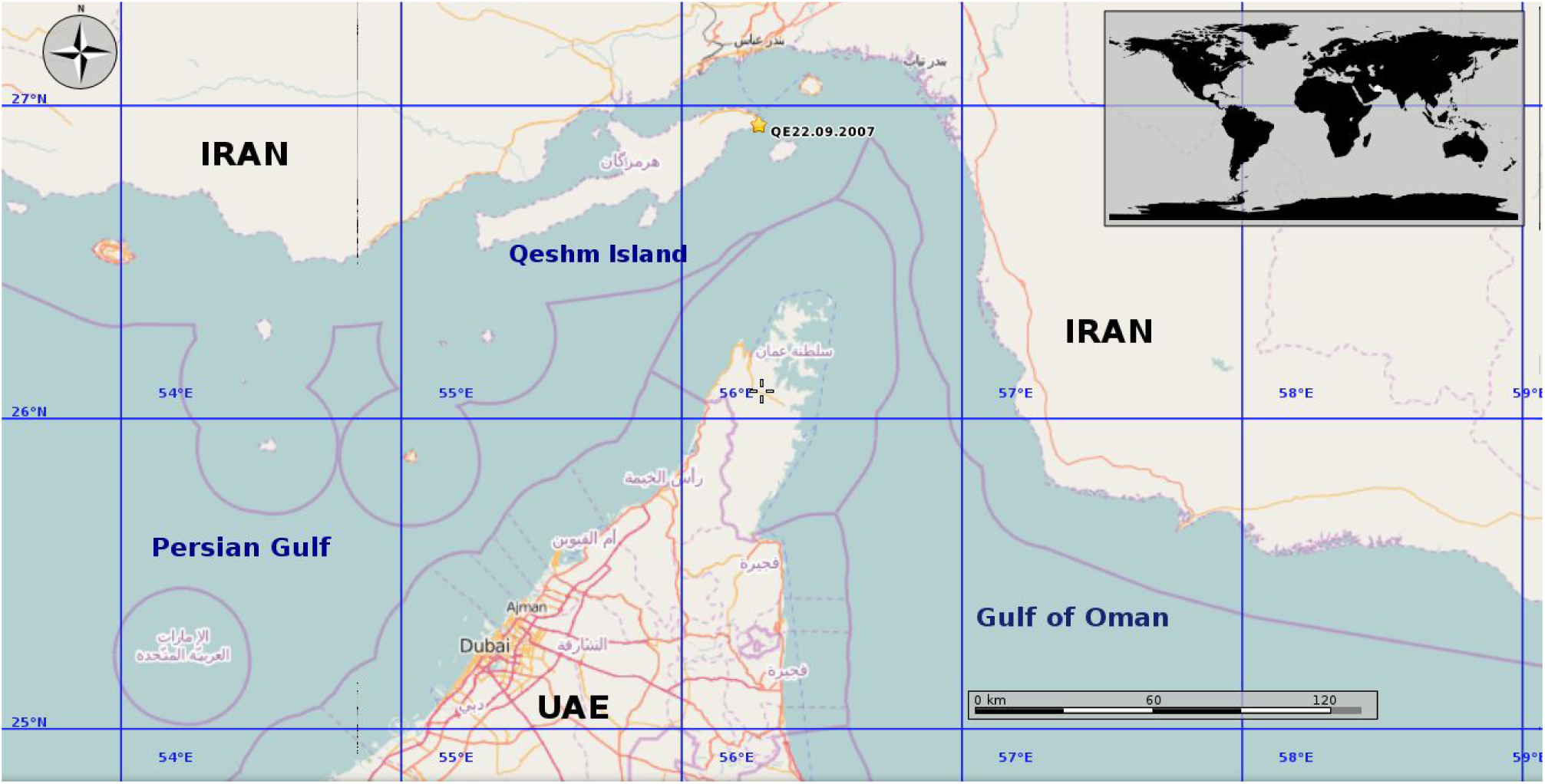
Map showing the stranding location of specimen QE22.09.2007 (yellow star) on the eastern tip of Qeshm Island, Iran, located at the Strait of Hormuz which links the Persian Gulf to the Gulf of Oman in the Northeast Indian Ocean.

## Results and Discussion

Below we discuss the morphologically diagnostic features which, one by one distinguish specimen QE22.09.2007 from some, most or all other balaenopterid species minus *B. omurai*.

### Diminutive body size

Body length of QE22.09.2007 measured 397 cm. Interestingly, a *B. omurai* calf, genetically identified as such, that stranded in Mauritania measured a virtually identical 398 cm (Jung et al., 2015; Mullié et al., 2015). The fully healed umbilicus of QE22.09.2007 (Figure 2) indicated an age class of, as a minimum, a several weeks-old neonate, but probably considerably older. QE22.09.2007 was significantly smaller than neonate length for blue whale (*B. musculus*), fin whale (*B. physalus*) and sei whale (*B. borealis*), which definitively discards the three larger balaenopterids from the differential diagnosis due to their large body size. In comparison, neonate size for Bryde’s whales ranges 3.81-3.96 cm (Best, 2007). Antarctic minke (*B. bonaerensis*) and common minke whales (*B. acutorostrata*), smaller still, range respectively 2.7-2.9 m (Best, 2007) and 2.0-2.8 m (Jefferson et al., 2008), species that cannot be excluded based on body size.

**Figure 2.**
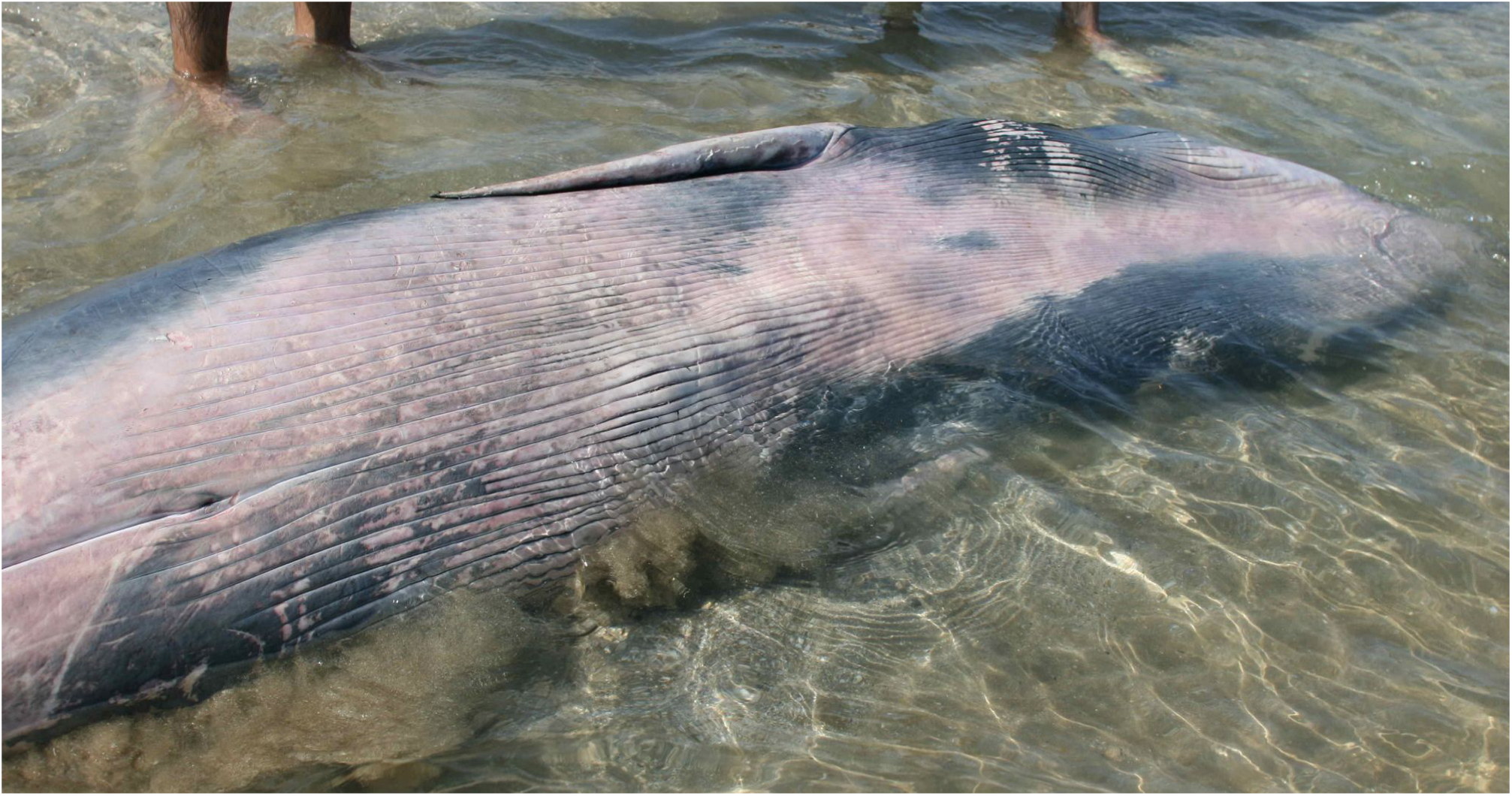
Ventral view of specimen QE22.09.2007, showing diagnostically large number of throat grooves (n=82), which extended caudad of the fully healed umbilicus. Note also the asymmetric ventral colouration pattern (see text).

### Large number of ventral grooves that extend caudad of umbilicus

From photographs it was determined that specimen QE22.09.2007 had about 82 narrow throat grooves which extended caudad of the umbilicus (Figure 2). This exceptionally large number is consistent with the 80-90 throat grooves reported for *B. omurai* (Wada et al., 2003) and the ‘minimum of 70’ in a juvenile specimen from Mauritania (Jung et al., 2015) that was similar to QE22.09.2007. Moreover, the lengthy throat grooves extended markedly caudad to the pectoral fin tips, as is evident also in the Mauritania individual (Jung et al., 2015).

Such a high number, and far caudad extension, of throat grooves distinguish QE22.09.2007 from *B. bonaerensis* which, on average, has merely some 65 grooves (range= 44-76) that extend in the midline from the tip of the lower jaw to just anteriad of the umbilicus (Best, 2007). These features distinguish QE22.09.2007 also from dwarf minke whale (*B. acutorostrata* subsp.) which has even less throat grooves, namely 44-66 at the cross-section of the flippers (Kato and Fujise, 2000) and grooves that extend only to anteriad of the umbilicus (Best, 2007). Ventral groove characteristics by itself exclude both minke whale species.

In Bryde’s whales the major ventral grooves number 42-54 and extend from the chin as far back as the umbilicus (Olsen, 1913; Best, 2007). Jefferson et al. (2008) indicated a higher upper range (40-70) for *B. brydei* but even that maximum does not compare positively with the 82 grooves observed in QE22.09.2007.

#### Strongly falcate dorsal fin

Omura’s whales have a strongly falcate and pointed dorsal fin (Wada et al., 2003; Jefferson et al., 2008; Cerchio et al., 2015), probably more consistently backswept than in any other rorqual species (Figure 3), especially in adults. Some growth variation may exist. Indeed, in several Omura’s whales, the often elongated tip points backwards at a 90° angle from vertical (e.g., photo on p.58, by T. Yamada *in* Jefferson et al., 2008; and supplementary material of Cerchio et al., 2015). This results in a very low dorsal fin height/length-at-base (H/L) ratio, 0.41 in the Yamada specimen. In QE22.09.2007, the dorsal fin was also strongly falcate and formed a 90° angle (Figure 3). Height and length at base measured, respectively, 7.5 cm and 18.0 cm, also resulting in a low H/L ratio of 0.42.

**Figure 3.**
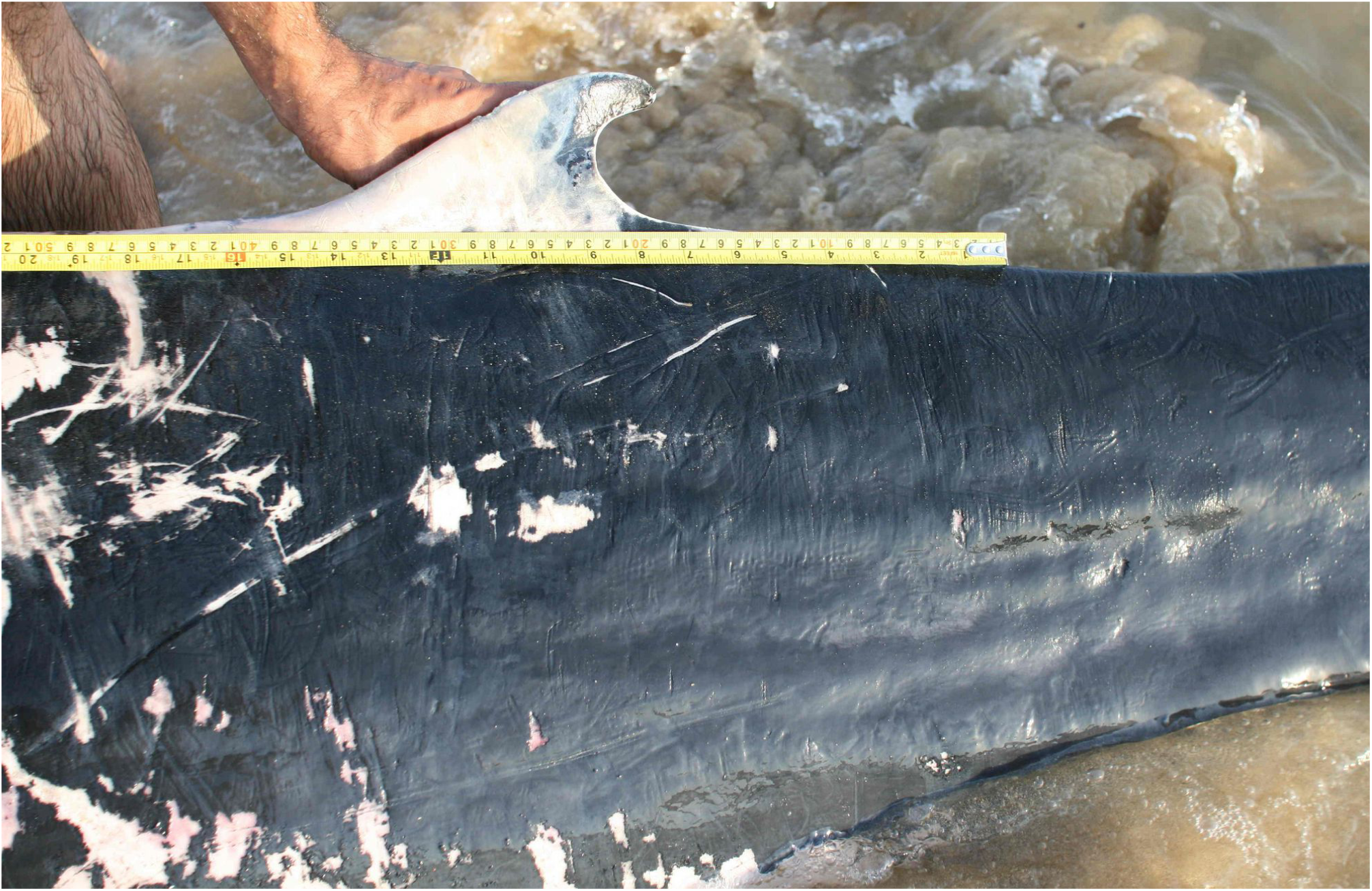
The strongly falcate, but relatively low, dorsal fin of QE22.09.2007, with tip pointing backwards seems a consistent characteristic for *B. omurai*. Indications are that the recurvedness accentuates with growth.

#### Asymmetric colouration pattern

Overall the colouration pattern of *B. omurai* resembles that of the fin whale *Balaenoptera physalus* (Wada et al. 2003; Jefferson et al., 2008). In QE22.09.2007 (Figure 2), as in *B. omurai*, the left side of the throat is darkly pigmented while much of the remaining ventral surface is mostly lightly coloured (Wada et al., 2003). However the lower lips, i.e. the skin covering the mandibula were a slightly darker grey (Figure 4), also clearly visible, albeit not discussed, in an adult female *B. omurai* from Madagascar (Cerchio et al., 2015, see first row of their Figure 3).

**Figure 4.**
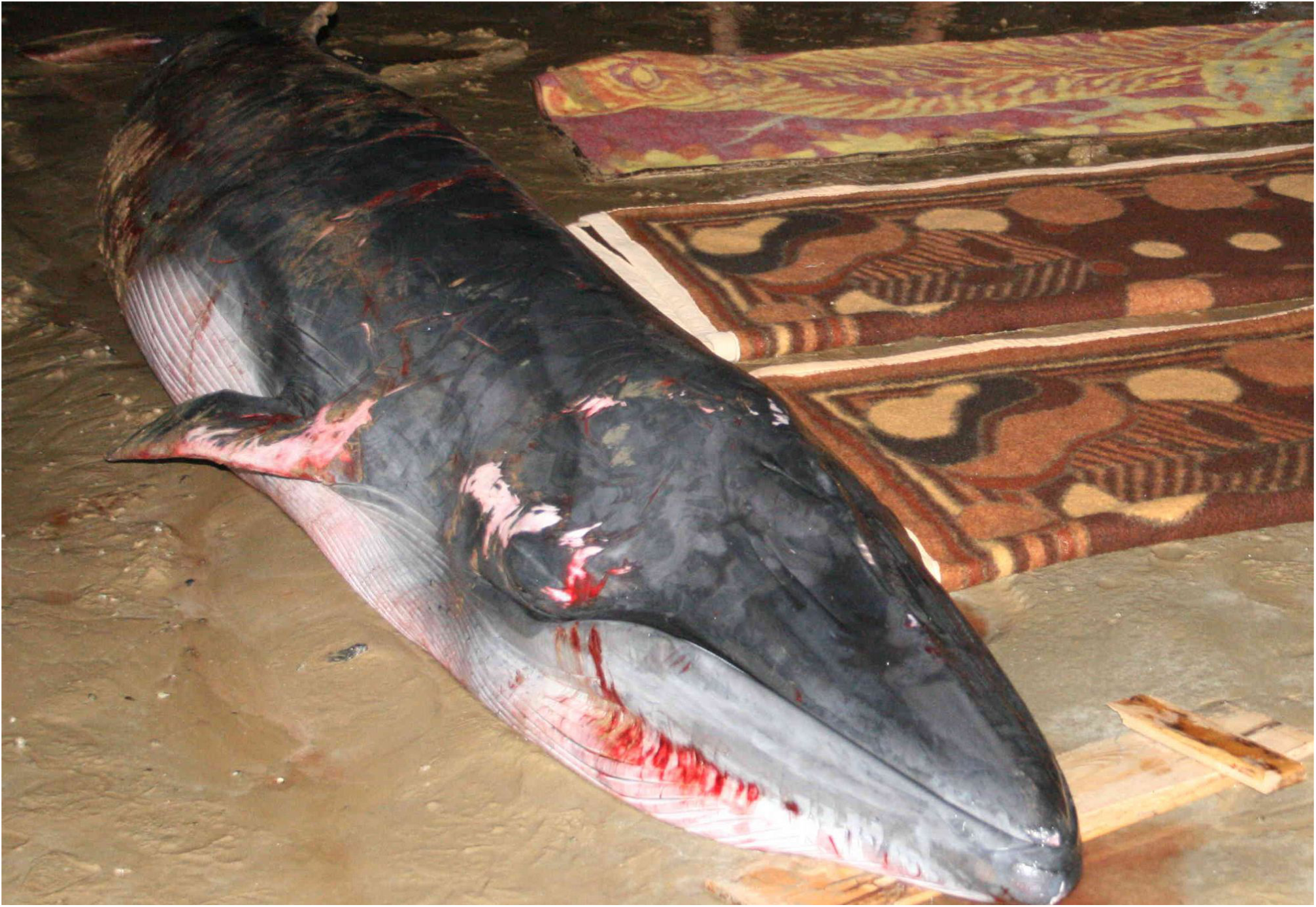
Right side of QE22.09.2007 while still alive, showing lightly pigmented, almost white throat, with light grey right mandible. Note three head stripes: (1) dark grey, wide eye stripe; (2) narrow ear stripe; and (3) wide flipper-to-flank stripe. Visible are also the central rostral ridge and two faint partial accessory ridges.

In summary, all features of colouration reported for *B. omurai* by Cerchio et al. (2015) were also evident in QE22.09.2007: (i) asymmetrical colouration of the lower jaws, with lightly pigmented right jaw (Figure 4) and darkly pigmented left jaw (Figure 5); (ii) asymmetrical colouration of the gape (inner lower lip) with whitish left gape (Figure 5) and darkly pigmented right gape; (iv) dark eye and ear stripes on the right side (Figure 4); (v) a third dark stripe (flipper-to-flank). A lightly pigmented chevron anterior to the dorsal fin and a blaze anterior to the right eye as reported by Cerchio et al. (2015) were not noticeable on photos of QE22.09.2007 and may, or may not, have been present. We must consider also the possibility of ontogenetic variation, as published descriptions are from subadults and adults, while QE22.09.2007 was evidently a calf. Whether the leading edge of the pectoral fins was white from tip to shoulder (Cerchio et al., 2015) neither could be ascertained, due to peeling skin in QE22.09.2007.

**Figure 5.**
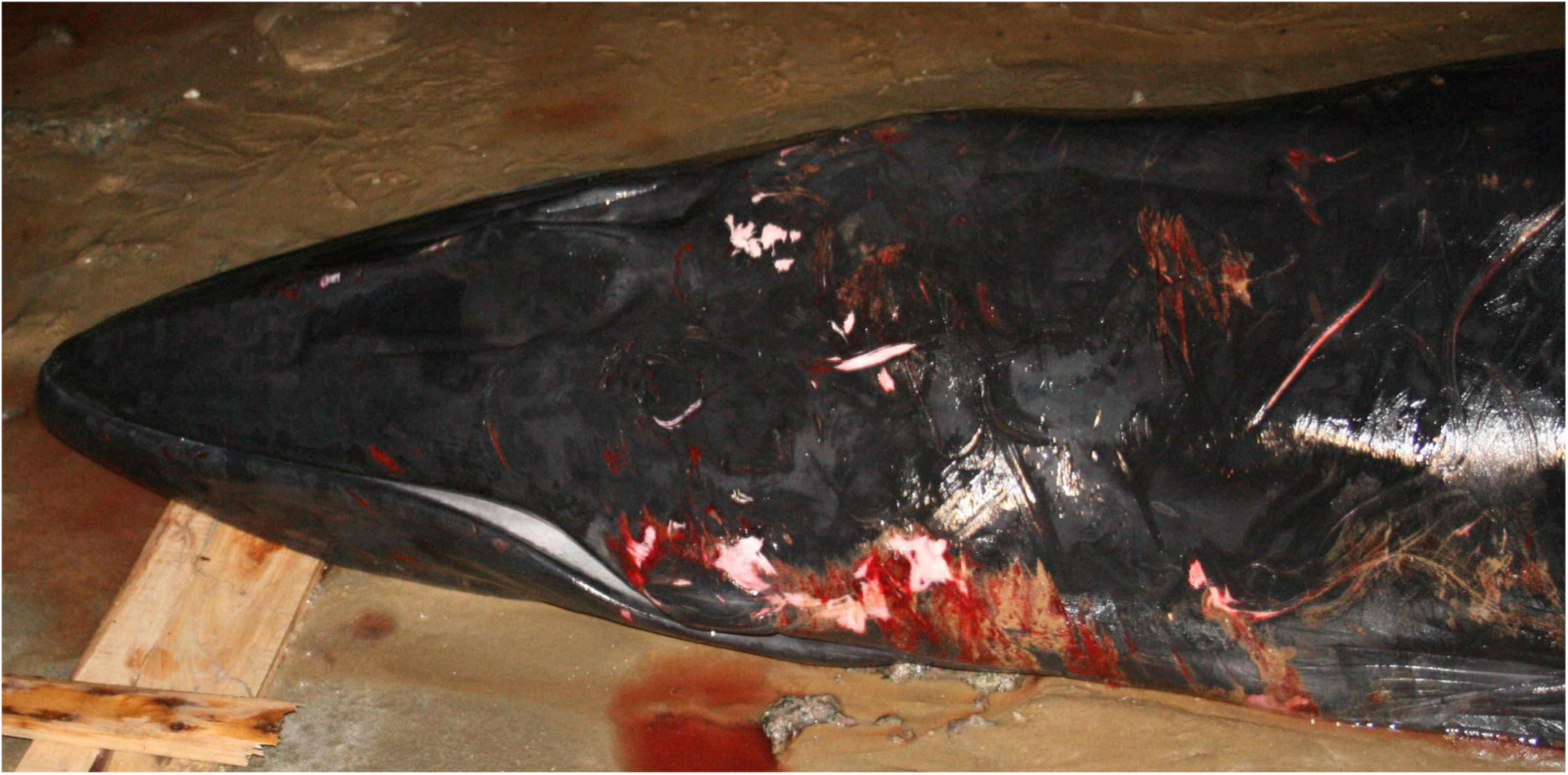
Overal dark pigmentation of the left body side of specimen QE22.09.2007, and in particular the uniformly darkly pigmented throat and left mandible contrasts with its paler right side (Figure 4). However note lighter pigmented left gape

#### Faint indications of lateral rostral ridges

In contrast with the prominent lateral ridges on the rostrum in Bryde’s whales, commonly characterized as diagnostic, the lateral rostral ridges are faint in Omura’s whales (Wade et al., 2003; see Figure 3 in Cerchio et al., 2015). In QE22.09.2007 a faint left lateral ridge was formed between two parallel grooves present on the left side of the rostrum, while on the right side, relief of the rostral integument was visible only proximally due to two very short parallel grooves (Figure 6). Actually these sets of parallel grooves appeared to be responsible for the faint lateral ridge aspect, as there seemed to be no true raised ridges as in Bryde’s whales. More *B. omurai* specimens will be necessary to better describe the extent of individual and ontogenetic variation in rostral grooves and ridges. However, it is evident that both minke whale species which have a single strongly raised central ridge on the rostrum, and laterally show no relief of the rostral integument, are not concordant with the morphology observed in QE22.09.2007.

**Figure 6.**
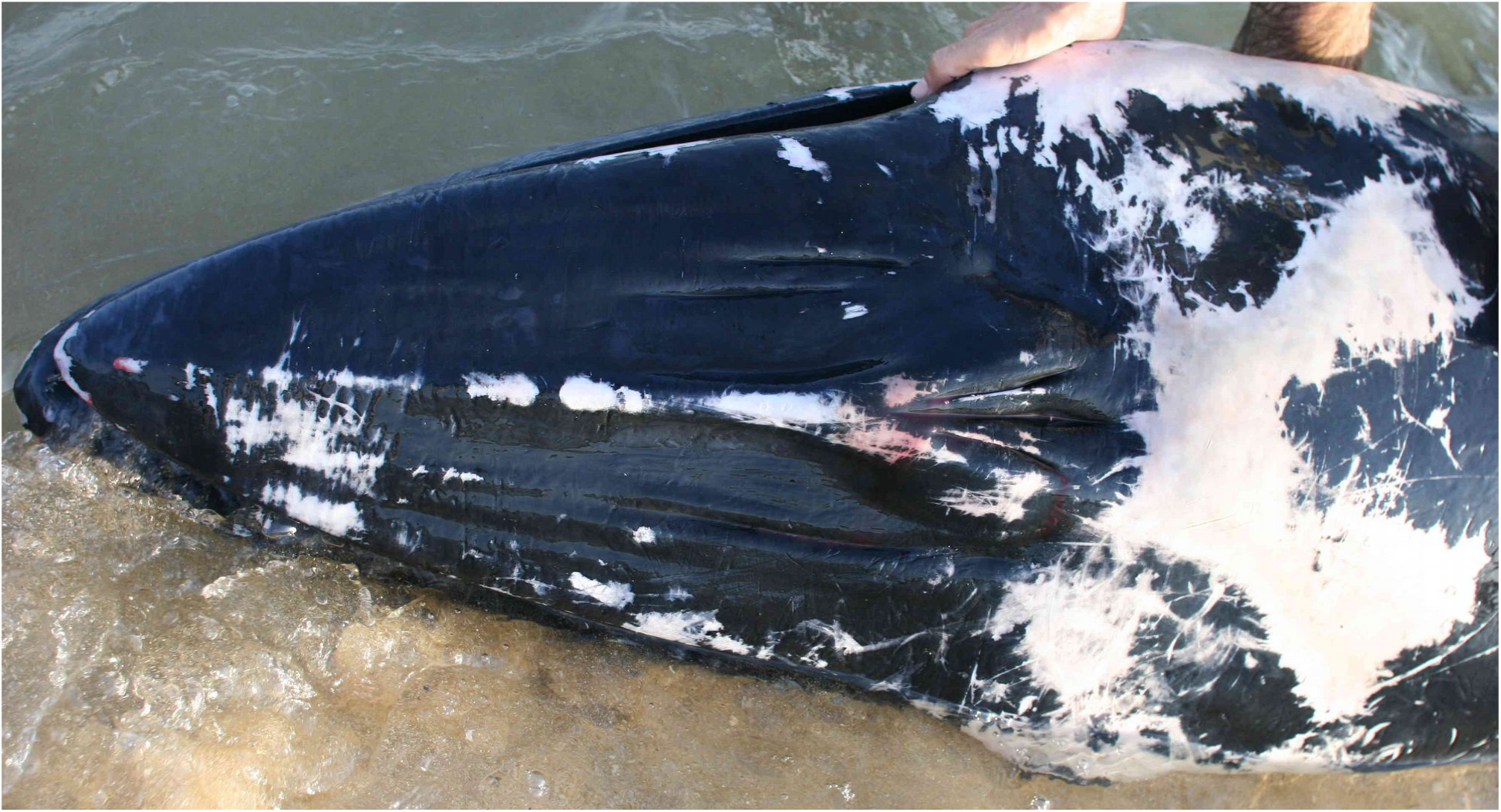
Dorsal view of head of QE22.09.2007 showing peculiar configuration of a relatively low central rostral ridge, a faint left lateral ridge and a system of two very short grooves, proximally, on the right side of the rostrum, providing a general aspect of incomplete and faint lateral ridges The rostrum was largely flat and not slightly arched in cross-section as in minke whales.

#### Low number of baleen plates

An approximate count of 204 baleen plates in the right jaw, determined from several close-up photographs of the head of QE22.09.2007 (Figure 7) agrees with baleen plate counts reported for *B. omurai*: 203 (right side) in one whale and 208 (left side) in another whale, while 181-190 (right side) in a third specimen (Wada et al., 2003). Baleen counts in *B. omurai* are markedly lower than those for the other *Balaenoptera* species (Wada et al., 2003). There exists no overlap even with the lower extreme of the range in Bryde’s whales (276-289 plates per jaw) (Best, 2007) and Antarctic minke whales (215-310 plates per jaw) (Best, 2007).

**Figure 7.**
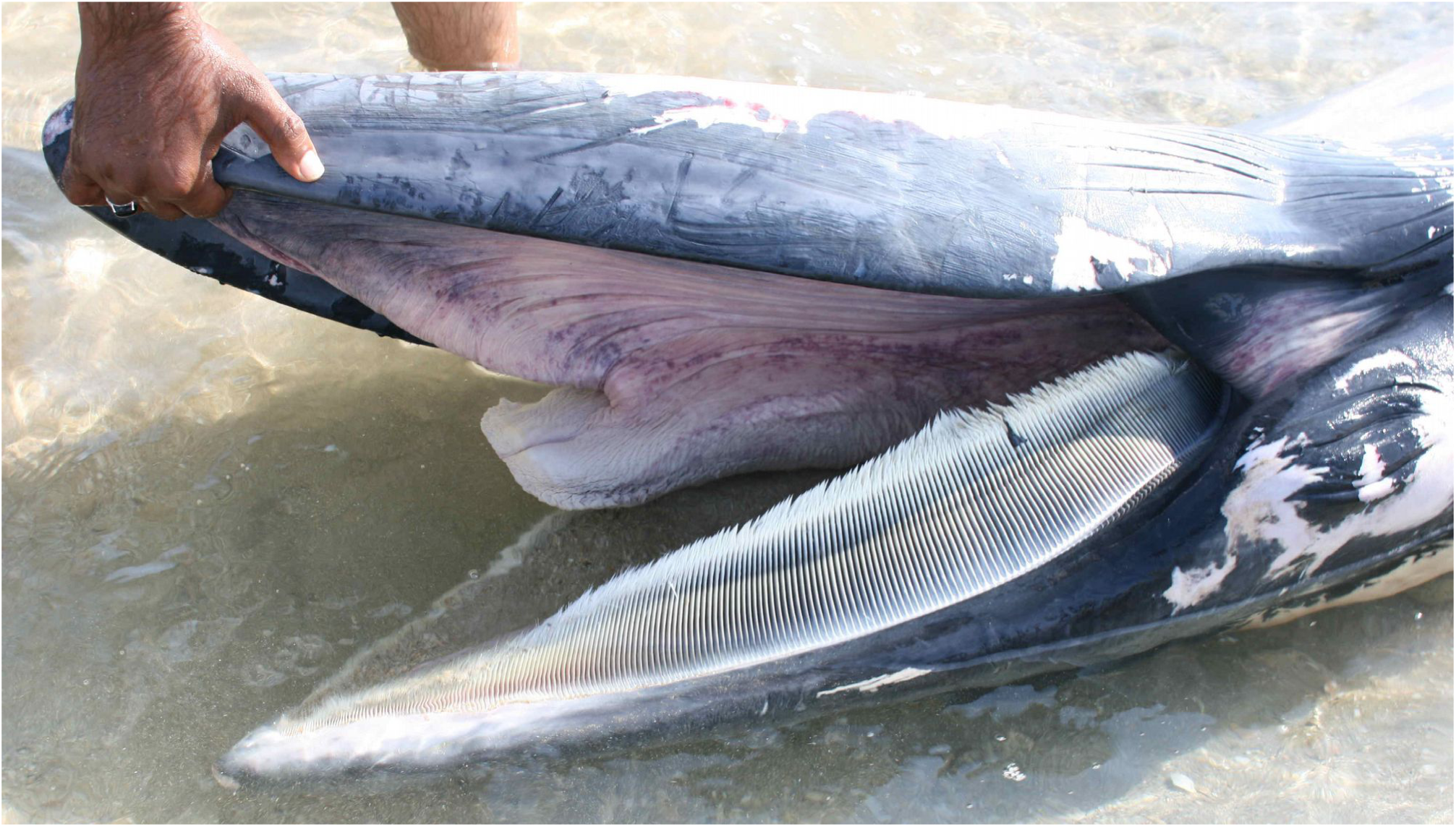
Right side view of the head (upside down) of specimen QE22.09.2007 showing the low number of short but broad baleen plates, with greyish-white baleen fringes, as reported for *B. omurai* (Wada et al., 2003). The colour of the baleen plates varied with their anteroposterior position in the baleen row and displayed asymmetry.

## Conclusion

The combination of seven principal morphological features is considered diagnostic evidence for the positive identification of QE22.09.2007 as *B. omurai*. With no evidence of emaciation or any external trauma other than minor cuts and abrasions associated to the event of running ashore, the cause of stranding is unknown. However, the event was strongly reminiscent of the recent stranding of a similar-sized juvenile specimen of *B. omurai* in Mauritania (Jung et al., 2015). As the latter, QE22.09.2007 apparently was also either an unweaned calf or a recently weaned one. An adult female (mother) may have occurred in the vicinity. The likelihood for a local *B. omurai* population in the northern Arabian Sea seems higher than the wandering of a very young animal or mother/calf pair from any of the known distant distribution areas in the eastern Indian Ocean or in the SW Indian Ocean (Madagascar).

Generally, Omura’s whale may be far more wide-spread than currently available records seem to suggest (reviewed in Cerchio et al., 2015). In the northeast Indian Ocean, there are specimens from the west coast of Thailand (Yamada et al., 2006; Adulyanukosol, 2012) and graphic evidence from the eastern Andaman Sea. The species occurs also on the west side of the Malay Peninsula, in Malaysia (Ponnanpalam, 2012) and off the Cocos Islands (Wada et al., 2003). However genetic sampling of Bryde’s whale populations in the North Indian Ocean, including the Arabian Sea, Bay of Bengal, the Maldives and south of Java, have not yet revealed evidence of *B. omurai*. The suggestion by Cerchio et al. (2015) that the species distribution may be discontinuous and that the Madagascar population in the southwest Indian Ocean may be relatively isolated from the eastern populations is possible but as yet unconfirmed. These authors admitted however that it is conceivable that other populations have gone undetected, with accounts from the Cocos Islands and Madagascar suggesting that populations occur around oceanic islands. Boat surveys should be undertaken east and south of Qeshm Island in search of a potential Iranian population of Omura’s whale.

It is adamant also that more stranded whales be systematically examined, photographed and sampled in the Persian Gulf. We know for instance that from March to October 2015 at least four whales stranded on Iran’s Persian Gulf coasts (S. Ranjbar, unpublished data), but none have been studied.

The successive findings of *B. omurai* in formerly unrecognised areas of distribution (Madagascar, Mauritania, Iran) hint that the more likely scenario is of a far wider tropical and subtropical, perhaps pantropical, distribution of *B. omurai* than hitherto documented. Earlier specimens may have been incorrectly identified as Bryde’s whales considering that most diagnostic morphological features of *B. omurai* have been described only in the past few years (Wada et al., 2003; Cerchio et al., 2015; Jung et al., 2015).

## Acknowledgements

The authors thank Mr Asghar Besharati for kindly providing a video copy for examination. Van Waerebeek warmly thanks the Qeshm Department of Environment, Qeshm Free Area, Qeshm City, Iran, for supporting two short study visits to Qeshm Island in 2014-2015.

